# Compendium of secondary metabolite biosynthetic diversity encoded in bacterial genomes

**DOI:** 10.1101/2021.08.11.455920

**Authors:** Athina Gavriilidou, Satria A. Kautsar, Nestor Zaburannyi, Daniel Krug, Rolf Müller, Marnix H. Medema, Nadine Ziemert

**Author notes:** These authors contributed equally: Athina Gavriilidou, Satria A. Kautsar. These authors jointly supervised this work: Marnix H. Medema, Nadine Ziemert.

## Abstract

Bacterial specialized metabolites are a proven source of antibiotics and cancer therapies, but whether we have sampled all the secondary metabolite chemical diversity of cultivated bacteria is not known. We analysed ~ 170,000 bacterial genomes and ~ 47,000 metagenome assembled genomes (MAGs) using a modified BiG-SLiCE and the new clust-o-matic algorithm. We found that only 3% of the natural products potentially encoded in bacterial genomes have been experimentally characterized. We show that the variation of secondary metabolite biosynthetic diversity drops significantly at the genus level, identifying it as an appropriate taxonomic rank for comparison. Equal comparison of genera based on Relative Evolutionary Distance revealed that *Streptomyces* bacteria encode the largest biosynthetic diversity by far, with *Amycolatopsis*, *Kutzneria* and *Micromonospora* also encoding substantial diversity. Finally we find that several less-well-studied taxa, such as Weeksellaceae (Bacteroidota), Myxococcaceae (Myxococcota), *Pleurocapsa* and Nostocaceae (Cyanobacteria), have potential to produce highly diverse sets of secondary metabolites that warrant further investigation.

## Introduction

Specialized metabolites (also called secondary metabolites) are biomolecules that are not essential for life but rather offer specific ecological or physiological advantages to their producers allowing them to thrive in particular niches. These Natural Products (NPs) are more chemically diverse than the molecules of primary metabolism, varying in both structure and mode of action among different organisms^1^. Historically, microbial NPs and their derivatives have contributed and continue to contribute a substantial part of chemical entities brought to the clinic, especially as anticancer compounds and antibiotics^2–4^. Regrettably, the emergence of antibiotic-resistant pathogens^3^ concomitant to a stagnation of antimicrobial discovery pipelines^2,4^ is leading to a global public health crisis^3^.

Nonetheless, genomics-based approaches to NP discovery^5,6^ have revealed a largely untapped and much more diverse source of biosynthetic potential within genomes^3,7^. These findings were possible following the discovery that bacterial genes encoding the biosynthesis of secondary metabolites are usually located in close proximity to each other, forming recognizable Biosynthetic Gene Clusters (BGCs). However, while the numbers and kinds of BGCs clearly differ across microbial genomes^7,8^ and metabolomic data indicate that some biosynthetic pathways are unique to specific taxa^9^, a systematic analysis of the taxonomic distribution of BGCs has not yet been performed. Similarly, while useful estimates of the chemical diversity of specific taxa have been provided^8^, methodical comparisons across taxa are lacking. Because of this, the scientific community appears undecided on the best strategy for natural product discovery: should the established known NP producers be studied further or should the community be investigating underexplored taxa^7,10^? A relatively recent question is how much chemical diversity is hidden in uncultured bacteria. Metagenomic assembled genomes from uncultured bacteria have demonstrated a big potential of unknown BGCs^7^. It is unclear to what extent unexplored associated ecological niches and (micro)environments are also associated with unique and unexplored chemistry.

Here, we harnessed recent advances in computational genomic analysis of BGCs to survey the enormous amount of genome data accumulated by the scientific community so far. Using a global approach based on more than 170,000 publicly available genomes, we created a comprehensive overview of the biosynthetic diversity found across the entire bacterial kingdom. We clustered 1,185,995 BGCs into 62,449 Gene Cluster Families (GCFs), and calibrated the granularity of the clustering to make it directly comparable to chemical classes as defined in NP Atlas^11^. This facilitated an analysis of the variance of diversity across major taxonomic ranks, which showed the genus rank to be the most appropriate to compare biosynthetic diversity across homogeneous groups. This finding allowed us to conduct comparisons within the bacterial kingdom. Evident patterns emerged from our analysis, revealing popular taxa as prominent sources of both actual and potential biosynthetic diversity, and multiple yet uncommon taxa as promising producers.

## Main text

### Biosynthetic diversity of the bacterial kingdom

To assess the global number of Gene Cluster Families found in sequenced bacterial strains, we ran AntiSMASH^12^ on ~170,000 genomes from the NCBI RefSeq database^13^ (Supplementary Table 1), spanning 48 bacterial phyla containing 464 families (according to the Genome Taxonomy DataBase classification - GTDB^14^). We also included almost 50,000 bacterial Metagenome Assembled Genomes (MAGs) from 6 metagenomic projects of various origins^15–20^ (Table 1 and Supplementary Table 1). To accurately group similar BGCs – which likely encode pathways towards the production of similar compounds – into Gene Cluster Families (GCFs) across such a large dataset, we used a slightly modified version of the BiG-SLiCE tool^21^, which has been calibrated to output GCFs that match the grouping of known compounds in the NP Atlas database^11^ (see Methods: Quantification of biosynthetic diversity with BiG-SLiCE). The resulting GCFs were then used to measure biosynthetic diversity across taxa.

**Table 1.** Input datasets and biosynthetic diversity with different BiG-SLiCE cut-offs. The “Complete Dataset” was used for the computation of the actual and potential biosynthetic diversity found in all cultured (and some uncultured) bacteria. The dataset “RefSeq bacteria with known species taxonomy” was used for pinpointing the emergence of biosynthetic diversity, for which accurate taxonomic information was needed, and for identifying groups of promising producers. The “T”s under Gene Cluster Families represent different BiG-SLiCE l2-normalized euclidean thresholds; the values under T=0.4 stand out due to it being considered the most suitable cut-off. BGC to GCF assignment for each threshold can be found in Supplementary Tables 2-5. *MAG sources: bovine rumen^15^, chicken caecum^16^, human gut^17^, ocean^18^, uncultivated bacteria^19^, various sources^20^.

| Dataset |  | Genomes | BGCs | Gene Cluster Families |  |  |  |
| --- | --- | --- | --- | --- | --- | --- | --- |
|  |  |  |  | T = 0.4 | T = 0.5 | T = 0.6 | T = 0.7 |
| Complete Dataset | All RefSeq bacteria | 170,549 | 1,060,592 | **51,052** | 37,785 | 28,057 | 19,152 |
|  | Bacterial MAGs* | 47,098 | 125,403 | **21,354** | - | - | - |
|  | Total | 217,647 | 1,185,995 | **62,449** | - | - | - |
| RefSeq bacteria with known species taxonomy | Complete Genomes | 16,004 | 94,904 | **16,984** | 13,546 | 10,399 | 7,151 |
|  | Draft Genomes | 147,265 | 913,642 | **37,123** | 27,748 | 20,638 | 14,016 |
|  | Total | 163,269 | 1,008,546 | **41,870** | 31,237 | 23,227 | 15,766 |

The number of GCFs in RefSeq ranged from 19,152 to 51,052 depending on the cut-off used by BiG-SLiCE (Table 1). While, as expected, the pure numbers of the analysis changed based on the l2-normalized euclidean threshold, the overall tendencies observed remained the same (Figure 1a, Supplementary Figure 1). The effect that the chosen threshold has on these results presented a challenge to our investigation, as previous estimations have also shown great heterogeneity when different thresholds were used^7,8^, precluding direct comparisons of their predictions. As each BGC can be considered a proxy for its encoded pathways and their products, differing thresholds will result in different degrees of granularity in the grouping of compound structures (Extended Data Figure 1). Nevertheless, linear relationships are not always applicable, as shown previously^22^, and a specific threshold will need to be set anyway to make comparisons possible. For this, we sought to directly relate the choice of our BGC clustering threshold to the clustering of their compound structures. NPAtlas, a database of known microbial small molecules, provides hierarchical clustering of the compound structures via Morgan fingerprinting and Dice similarity scoring^11^. As many as 947 compounds in NPAtlas are mapped to a known BGC in MIBiG repository^23^, giving us the opportunity to use them as an anchor for choosing our clustering threshold. After mapping the BiG-SLiCE groupings of known BGCs from the MIBiG to the compound clusters in NPAtlas (Supplementary Figure 2), we chose a threshold of 0.4, as it provided the most congruent agreements between the two groupings, with v-score=0.94 (out of 1.00) and ΔGCF=-17.

**Figure 1.**
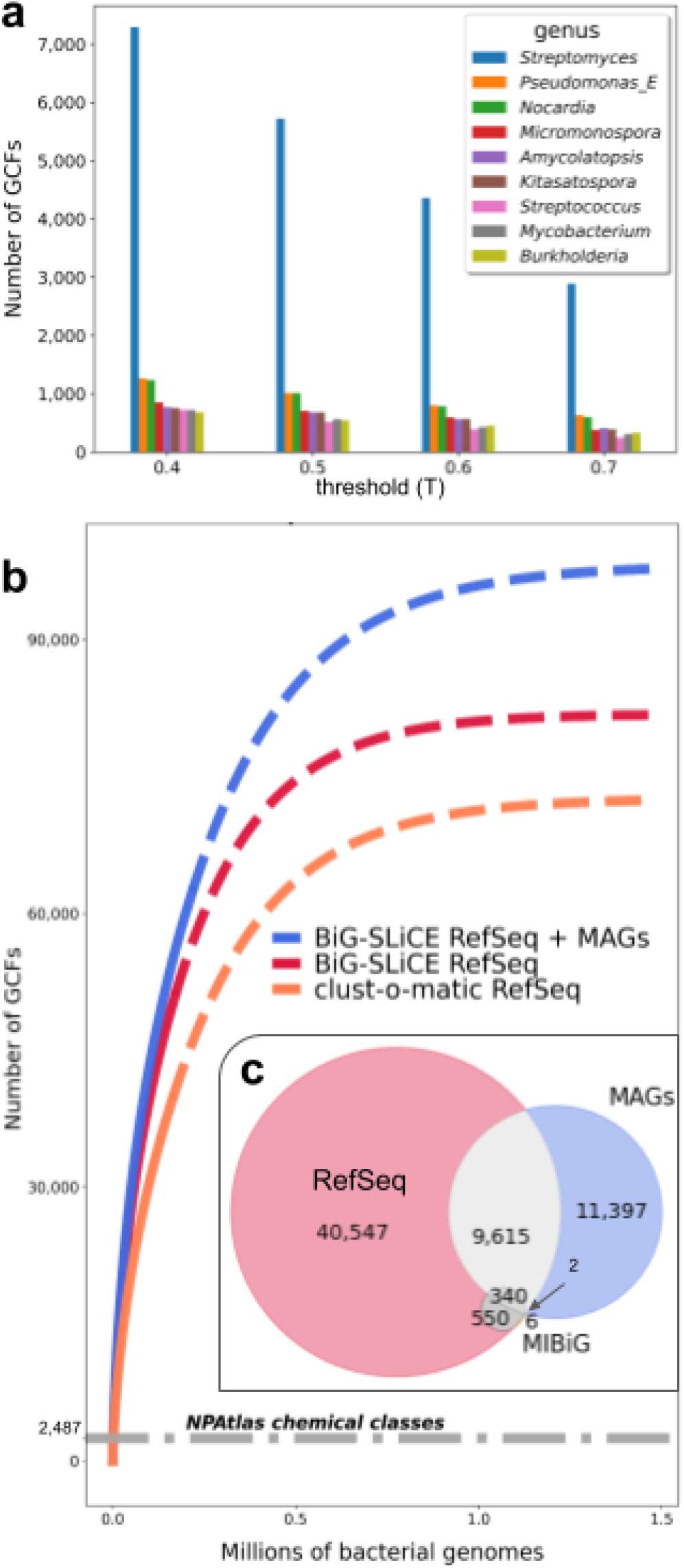
Biosynthetic diversity of the sequenced bacterial kingdom. Panel **a**: Bar plots of Gene Cluster Families (GCFs, as defined by BiG-SLiCE) of nine most biosynthetically diverse genera using different thresholds (T). The absolute number of GCFs changes from threshold to threshold, but the general tendencies (highest to lowest GCF count) are consistent between them. Panel **b**: Rarefaction curves of all RefSeq bacteria based on BiG-SLiCE (red) and based on clust-o-matic (orange), and rarefaction curve of the Complete Dataset, which includes bacterial MAGs (blue), based on BiG-SLiCE. BiG-SLiCE GCFs were calculated with T=0.4. Clust-o-matic GCFs were calculated with T=0.5. The solid lines represent interpolated and actual data, while the dotted lines represent extrapolated data. The number of chemical classes documented in NPAtlas^11^, which come from bacterial producers (gray dotted line - 2,487), corresponds to 2.5% - 3.3% of the predicted potential of the bacterial kingdom (number of GCFs at 1.6 million genomes). The Y values (number of extrapolated GCFs) at the right end of the graph are 97,760.12 (blue), 81,748.32 (red) and 72,411.11 (orange). Panel **c**: Venn Diagram of GCFs (as defined by BiG-SLiCE, T=0.4) of the bacterial RefSeq, Minimum Information about a Biosynthetic Gene cluster (MIBiG^23^) and bacterial MAGs datasets. More information on the MiBIG dataset can be found in Supplementary Table 6. About 53,4% of the GCFs of MAGs are unique (blue shape) to this dataset.

This calibration of thresholds of GCFs to families of chemical structures allowed us to perform a rarefaction analysis to assess how genomically encoded biochemical diversity (expressed as the number of distinct GCFs) increases with the number of sequenced and screened genomes (Figure 1b). The curve appears far from saturated, while the slope is steeper still if the bacterial MAGs are included in the analysis. When compared to the number of chemical classes documented in the NPAtlas^11^ database (Figure 1b), it appears that, to date, only about 3% of the kingdom’s biosynthetic diversity has been experimentally accessed.

In an attempt to evaluate the potential contribution of metagenomic data to Natural Product (NP) discovery, we studied how many of the GCFs found in the MAGs datasets were unique to this dataset (Figure 1c). Around 53,4% of GCFs in the MAGs were not found in the RefSeq strains or in the Minimum Information about a Biosynthetic Gene cluster database (MIBiG^23^). Paradoxically, in Figure 1b, the contribution of MAGs does not reflect this finding, but this is most likely because the metagenomic dataset is of limited size and does not cover the full microbial diversity of the biosphere. An analysis of the uniqueness of GCFs found in different environments, although only limited to one^20^ of the MAGs datasets, suggests that a connection exists between the biogeography of microbiomes and the uniqueness of their biosynthetic diversity, as the majority of GCFs (74.43 %) are biome-specific (Extended Data Figure 2, Supplementary Table 7).The latter finding is concordant with recent proof that most genes have a strong biogeography signal^24^.

### Variation in biosynthetic diversity drops at genus level

To identify the most promising bacterial producers, it is important to compare them at a specific taxonomic level. Several studies indicate that there is significant discontinuity in how BGCs are distributed across taxonomy: ‘lower’ taxonomic ranks like species within a genus carry more similar biosynthetic diversity, than ‘higher’ taxonomic ranks like phyla within a kingdom. To assess which taxonomic rank is the most appropriate to evaluate biosynthetic potential, we aimed to determine up to which taxonomic level the biosynthetic diversity remains homogeneous within that taxon. For this analysis, from our initial dataset, we left out the MAGs and only used the RefSeq bacterial strains as taxonomic assignment up to species rank (based on GTDB^14^) was available only for the latter dataset(Table 1).

We first decorated the GTDB^14^ bacterial tree with GCF values from the BiG-SLiCE analysis (Figure 2a), revealing the biosynthetic diversity found within currently sequenced genomes at the phylum rank. It immediately stood out that biosynthetic diversity was differently dispersed among the bacterial phyla, in accordance with published data^7,25^. As expected for known NP producers, the phyla Proteobacteria and Actinobacteria appeared particularly diverse^8,26,27^. However, these phyla are amongst the most studied and therefore the most sequenced^8,26,27^, a bias that was addressed later in the study.

**Figure 2.**
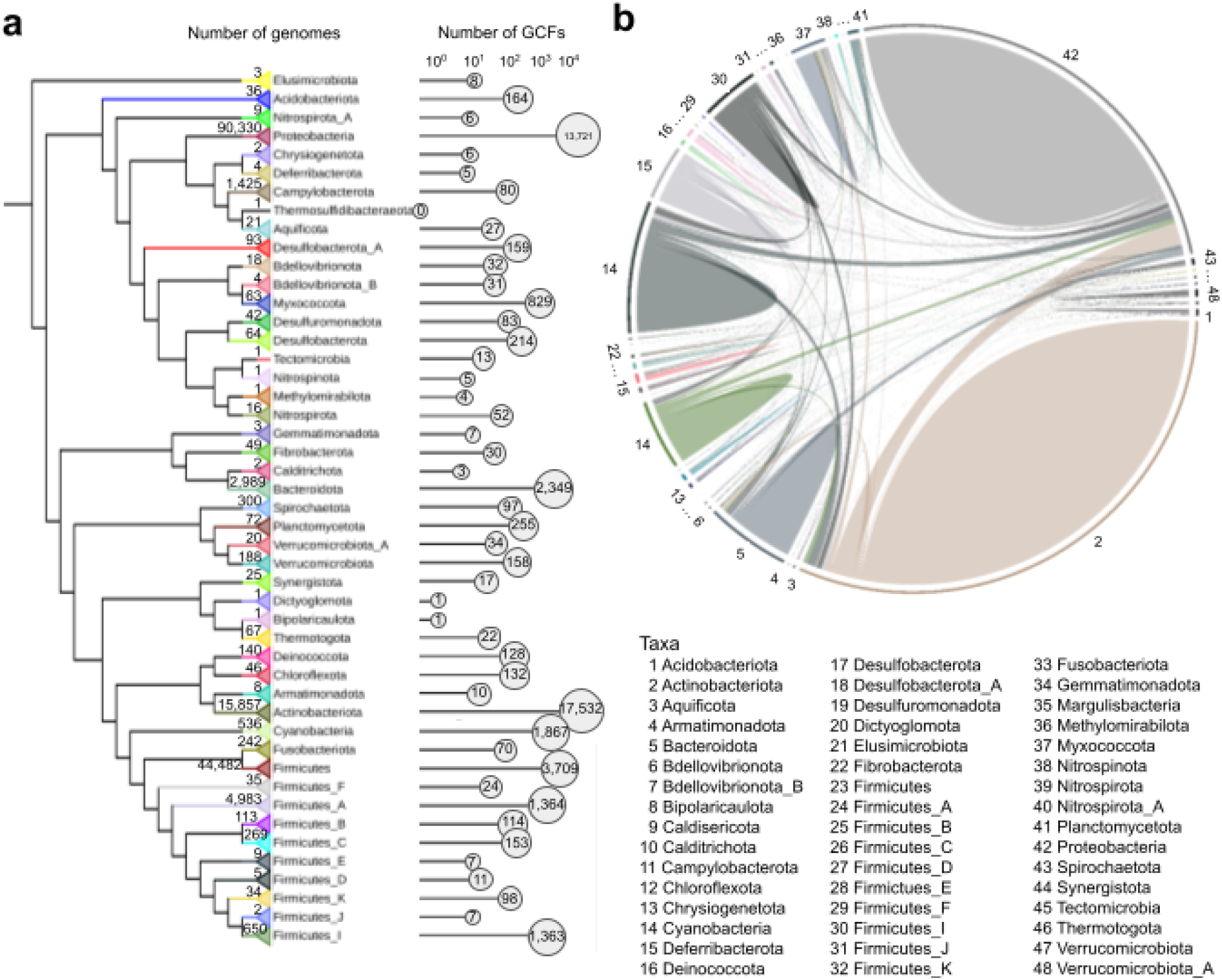
Comparison of biosynthetic diversity among phyla. Panel **a**: The Genome Taxonomy DataBase (GTDB^14^) bacterial tree was visualized with iTOL^30^ v6.5.2, decorated with Gene Cluster Families (GCFs) values (as defined by BiG-SLiCE at T=0.4), collapsed at the phylum rank and accompanied by bar plot of GCFs in logarithmic scale (10^0^ to 10^4^). The number of genomes belonging to each phylum is displayed next to the tree’s leaf nodes. Panel **b**: GCFs, as defined by BiG-SLiCE (T=0.4), unique to phyla (solid shapes) and with pairwise overlaps between phyla (ribbons), visualized with circlize^31^. Each phylum has a distinct color. Actinobacteriota (2) and Proteobacteria (40) seem particularly rich in unique GCFs.

Next, we examined whether the diversity of each phylum contributed to the domain’s total diversity, or if there was overlap among them. For this reason, we depicted the number of unique GCFs within each phylum, as well as the pairwise overlaps (Figure 2b). In most phyla, the vast majority (on average 73.81 ± 20.35%) of their GCFs appeared to be unique to them and not found anywhere else. This is coherent with the fact that HGT events, although relatively frequent for BGCs^28^, are much more common among closely related taxa^29^.

Once we obtained information on the diversity of different phyla, as well as the rest of the major taxonomic ranks (classes, orders, families, genera, species), we proceeded to determine from which taxonomic rank biosynthetic diversity levels no longer show high variability. Therefore, we conducted a variance analysis that included each taxonomic rank, from phylum to species. For each rank, the variance value was computed based on the #GCFs values of immediately lower-ranked taxa (see Methods: Variance Analysis). The distribution of these variance values for each rank is visualized in Figure 3a.

**Figure 3.**
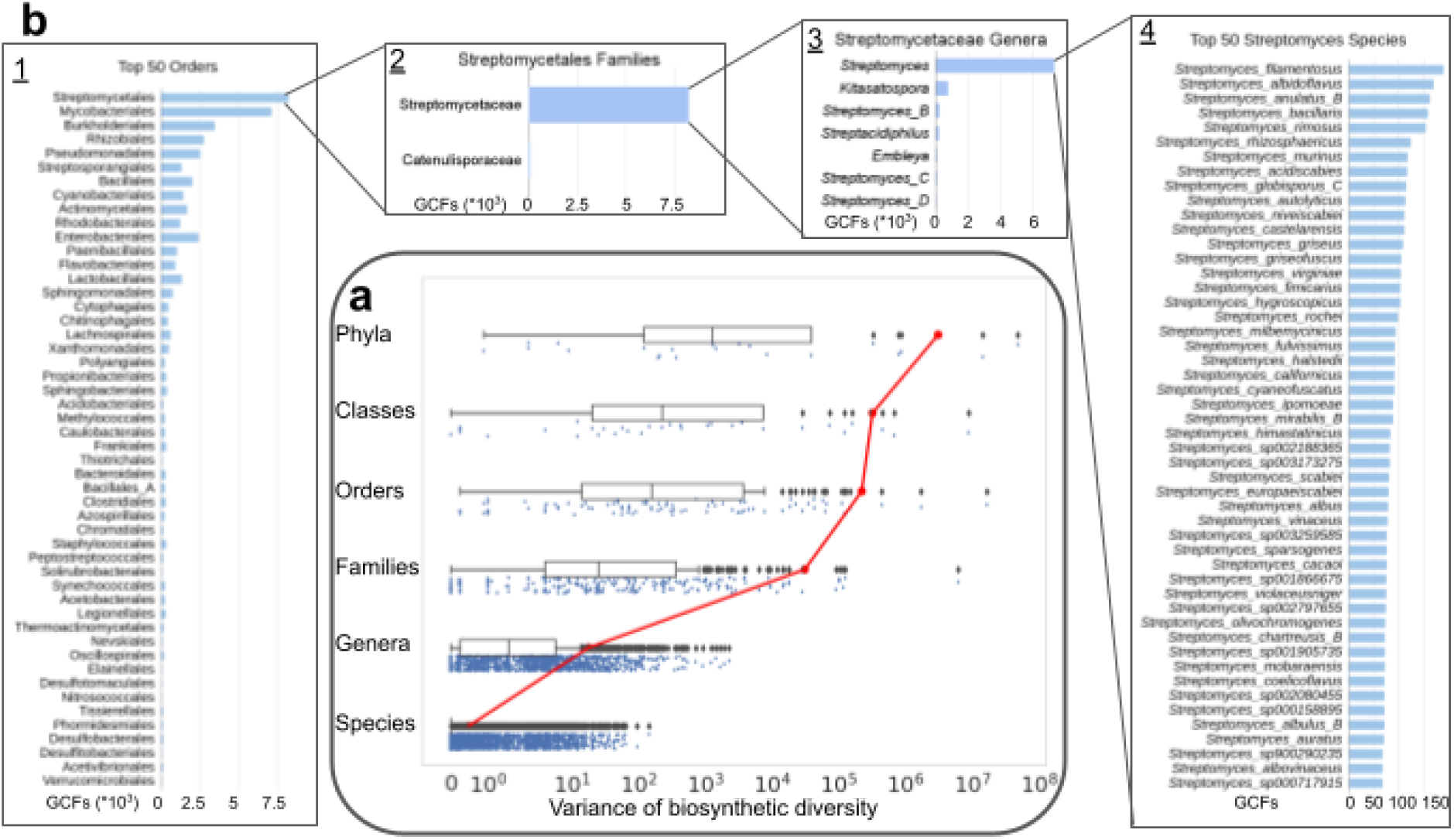
Relations of taxonomic levels to variability in biosynthetic diversity. Panel **a**: Modified “raincloud plots”^32^ of major taxonomic ranks (X axis in logarithmic scale). Each boxplot represents the dispersion of variance values of a certain taxonomic rank, computed from the number of Gene Cluster Families (GCFs as defined by BiG-SLiCE at T=0.4) of the immediately lower rank. The boxplots’ center line represents the median value; the box limits represent the upper and lower quartiles. Whiskers represent a 1.5x interquartile range. Points outside of the whiskers are outliers. Sample sizes are: Phyla n=21, Classes n=33, Orders n=89, Families n=224, Genera n=1,607, Species n=13,065. Jittered raw data points are plotted under the boxplots for better visualization of the values’ distribution. The red line connects the mean variance values of each rank. There is a noticeable drop in dispersion of variance values from the family rank to the genus rank (see also Supplementary Figure 3), indicating that the genera are suitable taxonomic groups to be characterised as diverse and be compared to each other. Panel **b**: Biosynthetic diversity of various taxa, measured in absolute numbers of distinct GCFs as defined by BiG-SLiCE (T=0.4) from currently sequenced genomes. Top 50 most diverse orders (<u>1</u>), Streptomycetales families (<u>2</u>), Streptomycetaceae genera (<u>3</u>), top 50 most diverse *Streptomyces* species (<u>4</u>). The difference in variance is visible in the graphs <u>1</u>,<u>2</u>,<u>3</u>, but becomes homogeneous at the species level as is shown in graph <u>4</u>.

There is a noticeable drop in the range of variance values for each rank, while diversity becomes highly homogeneous at the species level (Figures 3a,b). The plunge is most striking from the family to the genus level (Figure 3a), with even the outliers all falling under the 10^3^-line in the genus rank. Different species within a genus are likely to display uniform biosynthetic diversity, while much dissimilarity is observed between different genera belonging to the same family (Figure 3b). Additional statistical analysis confirmed the significance of this observation (Supplementary Figure 3) thus pinpointing, for the first time, the genus rank as the most appropriate for comparative analyses.

### Taxa that are sources of substantial biosynthetic diversity

The identification of the genus level as the most informative rank to measure biosynthetic diversity across taxonomy paved the way for a comprehensive comparative analysis of biosynthetic potential across the bacterial tree of life. However, to be able to systematically compare diversity values among groups, said groups need to be uniform. In this case, a common phylogenetic metric was necessary. We chose Relative Evolutionary Divergence (RED) and a specific threshold that was based on the GTDB’s range of RED values for the genus rank^14^ to define REDgroups: groups of bacteria analogous to genera but characterized by equal evolutionary distance (see Methods: Definition of REDgroups). Our classification revealed the inequalities in within-taxon phylogenetic similarities among the genera, with some being divided into multiple REDgroups (for example the *Streptomyces* genus was split into 21 REDgroups: Streptomyces_RG1, Streptomyces_RG2 etc.) and some being joined together with other genera to form mixed REDgroups (for example Burkholderiaceae_mixed_RG1 includes the genera *Paraburkholderia*, *Paraburkholderia_A*, *Paraburkholderia_B*, *Burkholderia*, *Paraburkholderia_E* and *Caballeronia*). This disparity among the genera reaffirmed the importance of defining the REDgroups as a technique that allowed for fair comparisons among bacterial producers.

The resulting 3,779 REDgroups showed huge differences in biosynthetic diversity as measured by the numbers of GCFs found in genomes sequenced from these groups so far, with the maximum diversity at 3,339 GCFs, average at 17 GCFs and minimum at 1 GCF. Nevertheless, the variance of diversity within the REDgroups was even more uniform than in the genera (Supplementary Figure 4). Some of the top groups (Extended Data Table 1, Supplementary Table 8) included known rich NP producers, such as *Streptomyces*, *Pseudomonas_E* and *Nocardia^23,26,27,33^*.

Although very informative, this analysis is biased because of large differences in the number of sequenced strains among the groups, with the economically or medically important strains having been sequenced more systematically than others. To overcome this bias, rarefaction analyses were conducted for each REDgroup (Figure 4b, Supplementary Table 8), as performed in previous studies^34,35^. Additionally, to examine how effectively this method overcomes the sequencing bias, a random sampling approach was taken (see Methods: Random sampling), which showed comparable results to the original analysis (Supplementary Table 9). With all the information on REDgroups, and in order to provide a global overview of the actual biosynthetic diversity and the potential number of GCFs, we modified and complemented the bacterial tree from Parks *et. al.^14^*, as shown in Figure 4a (Extended Data Figure 3). The dispersion of these values across the various phyla can also be seen, with the exceptional outliers standing out: Streptomyces_RG1, Streptomyces_RG2, Amycolatopsis_RG1, Kutzneria, and Micromonospora. All these are groups known for their NP producers^8,26,27,36^ and they remain in the top (Extended Data Table 1, Supplementary Table 8), seemingly having much unexplored biosynthetic potential.

**Figure 4.**
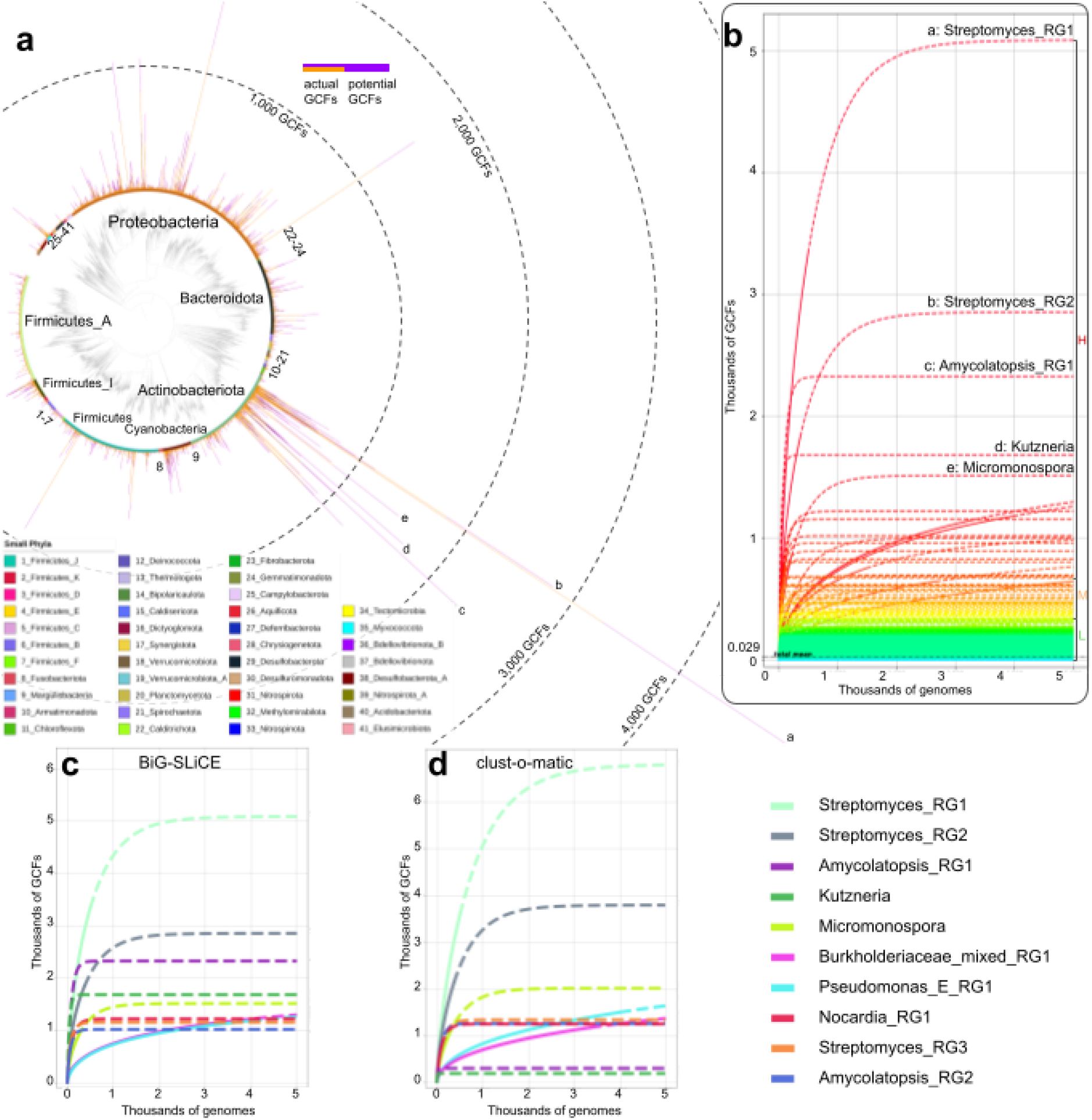
Overview of actual and potential biosynthetic diversity of bacterial kingdom, compared at REDgroup level. Panel **a:** GTDB^14^ bacterial tree up to REDgroup level, visualized with iTOL^30^ v6.5.2, colour coded by phylum, decorated with barplots of actual (orange) and potential (purple) Gene Cluster Families (GCFs), as defined by BiG-SLiCE (T=0.4). Top REDgroups with most potential GCFs include the following: A: Streptomyces_RG1, B: Streptomyces_RG2, C: Amycolatopsis_RG1, D: Kutzneria, E: Pseudomonas_E. Phyla known to be enriched in NP producers are immediately visible (Actinobacteriota, Protobacteriota), with the most promising groups coming from the Actinobacteriota phylum (the highest peak belongs to a REDgroup containing *Streptomyces* strains). Simultaneously, within the underexplored phyla, there seems to be significant biosynthetic diversity and potential. An interactive version of Figure 4a can be accessed online (Extended Data Figure 3). Panel **b:** Rarefaction curves of REDgroups (BiG-SLiCE T=0.4). In panels b, c and d the solid lines represent interpolated and actual data, while the dotted lines represent extrapolated data. The letters “L”, “M” and “H” correspond to Low- (0-389 pGCFs), Medium- (390-649 pGCFs) and High-diversity (more than 650 pGCFs) producers. The “L” range includes 3,737 REDgroups (shades of green), the “M” range includes 22 (shades of yellow/orange), while the “H” range includes 20 REDgroups (shades of red). The vast majority of REDgroups belong to the low-diversity producers (the mean of all REDgroups’ pGCFs is 29). The labels of most promising REDgroups are indicated (the letters a-e correspond to the peaks in panel a). *Streptomyces* strains are included in several of them. Panel **c:** Rarefaction curves of the most promising REDgroups (BiG-SLiCE T=0.4). Panel **d:** Rarefaction curves of the most promising REDgroups (clust-o-matic T=0.5). Though the exact numbers differ, the similarities between the two methods are apparent.

To ensure that our conclusions are not the product of algorithmic artifacts, we reran the analysis using an alternative method of quantifying biosynthetic diversity, which was developed independently, yet for the same purpose. This alternative approach, called clust-o-matic, is based on a sequence similarity all-versus-all distance matrix of BGCs and subsequent agglomerative hierarchical clustering in order to form GCFs (see Methods: Quantification of biosynthetic diversity with clust-o-matic). Like for BiG-SLiCE, we calibrated the threshold for clust-o-matic based on NP Atlas clusters. When comparing the results (Figure 4c,d, Supplementary Table 8), despite slight differences in absolute numbers, the two algorithms appeared to identify very similar trends.

*Streptomyces*, even when split into multiple REDgroups, is in the top groups both based on the known biosynthetic diversity and based on the estimated potential values. 5,908 (+103 Streptomyces_B, +39 Streptomyces_C, +16 Streptomyces_D) GCFs appear to be unique to the group, even among other phyla (Figure 5a). This is in agreement with previous studies investigating how much overlap there is among the main groups of producers^37^. What is more, streptomycetes appear to be the source of a good percentage of the biosynthetic diversity attributed to the Actinobacteria phylum, as seen in Figure 5b.

**Figure 5.**
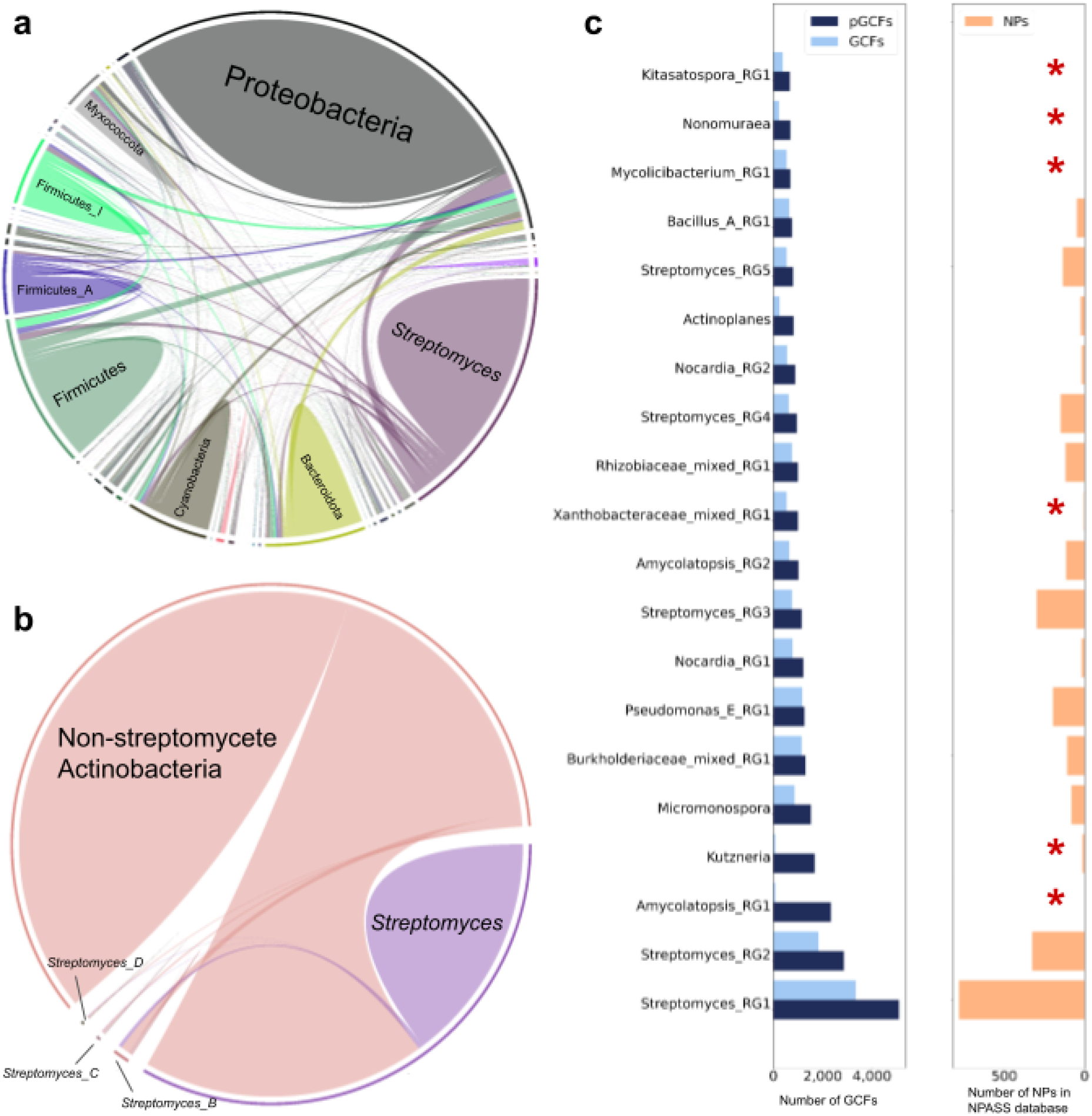
Unique diversity in the known producer *Streptomyces* and promising potential of less popular taxa. Panel **a**: Unique Gene Cluster Families (GCFs) as defined by BiG-SLiCE (T=0.4), of phyla and *Streptomyces* (solid shapes) and pairwise overlaps of phyla - phyla and phyla - *Streptomyces* (ribbons), visualized with circlize^31^. Each taxon has a distinct color. The smaller shapes and ribbons represent smaller phyla that can be seen in Extended Data Figure 4. The genus *Streptomyces* appears to have a very high amount of unique GCFs comparable to entire phyla, such as Proteobacteria. Panel **b**: Unique GCFs as defined by BiG-SLiCE (T=0.4), of non-streptomycete Actinobacteriota and all *Streptomyces* genera (solid shapes) and pairwise overlaps between Actinobacteriota and *Streptomyces* (ribbons), visualized with circlize^31^. The *Streptomyces* genus, only one of many belonging to the Actinobacteriota phylum, appears to be responsible for a big percentage of the phylum’s unique diversity (see big pink ribbon). Panel **c**: Left: Potential (pGCFs) and actual (GCFs) number of Gene Cluster Families as defined by BiG-SLiCE (T=0.4), of top 20 most promising REDgroups. Right: number of Natural Products (NPs) found in the NPASS database^38^, that originate from species included in each REDgroup. The REDgroups with few (< 15) to no known NPs associated with them are marked with red stars on the right side of the graph. Several of the displayed groups are in the latter category: Amycolatopsis_RG1, Kutzneria, Xanthobacteraceae_mixed_RG1 (containing the genera *Bradyrhizobium, Rhodopseudomonas, Tardiphaga* and *Nitrobacter*), Mycolicibacterium_RG1, Nonomuraea, Kitasatospora_RG1.

However, taxa less popular for NP discovery also show promise, as was evidenced by a comparison of our results with data from the NPASS database of Natural Products^38^ (Figure 5c). Among the 20 overall most promising REDgroups we found at least 6 groups that show promise but whose members are either not catalogued in the database as NP sources or are connected to few (<15) known compounds: Amycolatopsis_RG1, Kutzneria, Xanthobacteriaceae_mixed_RG1, Mycolicibacterium_RG1, Nonomuraea, Kitasatospora_RG1. The Amycolatopsis_RG1 group only includes three rare species: *Amycolatopsis antarctica, marina* and *nigrescens*. Other promising REDgroups with very few known producers include Cupriavidus (from Proteobacteria phylum), Weeksellaceae_mixed_RG1 (from Bacteroidota phylum) and Pleurocapsa (from Cyanobacteria phylum). More information about the promising underexplored taxa can be found in Supplementary Table 8.

## Discussion

Using two different algorithms, we mined deposited bacterial sequencing data to identify Biosynthetic Gene Clusters (BGCs) and grouped them into gene cluster families (GCFs) according to chemical families of encoded compounds. We identified maximal emergence of the highest biosynthetic diversity close to the genus rank and chose to further investigate analogous taxonomic groups (REDgroups). Rarefaction analysis identified the highest biosynthetic potential and the most promising bacterial taxa among many known diverse groups as well as multiple promising understudied producers. To the best of our knowledge, this is the largest survey of secondary metabolite production to date, and our study provides a reproducible pipeline to underpin drug discovery efforts.

The biosynthetic capacity of the bacterial kingdom was previously assessed by Cimermancic *et. al.^7^*, but the dataset analysed was 33,000 BGCs compared with the 1,185,995 BGCs we analysed. Additionally they used ClusterFinder, which is known as a more exploratory identification tool^7,39^. Projects that exploit publicly available genomic data are reliant on the quality of genomes sequenced as well as the efficiency of available genome mining methods, which have some limitations^40^. For instance, the study of GCF uniqueness among taxa may be affected by antiSMASH’s imperfect BGC boundary prediction^12^. Even though BiG-SLiCE converts BGCs into features based only on domains related to biosynthesis^21^, genomic context unrelated to the biosynthetic pathway of a BGC could still have a role in the GCF assignment; this issue cannot be fully addressed with currently available tools. However, antiSMASH’s ability to discern cluster limits and detect BGCs from cultured strains and MAGs is comparable to alternative tools, while its ability to predict different BGC types is unparalleled^41^, as is apparent from its common use in Natural Product (NP) research^7,9,25,33,35,42^. What is more, the fact that it is rule-based^12^ implies the possibility of undetected types of clusters and increases the likelihood that our calculations have underestimated the true biosynthetic potential of bacterial organisms.

Furthermore, our pipeline was the first to use the GTDB^14^ taxonomy for studying global bacterial biosynthetic diversity. This enabled us to avoid misclassifications of NCBI taxonomic placement^43–46^. The use of rarefaction curves allowed us to infer the biosynthetic potential of bacterial groups, as done in some smaller-scaled projects^7,8,34,35^. This method aims to enable fair comparisons among incomplete samples^47^. However, while overestimation is not expected to happen, for those groups that contain very few genomes, there is a tendency to underestimate their potential capacity^47^. Hence, sequencing bias of popular taxa still affects our results. We tried to minimize the bias within the pipeline as much as possible while retaining high diversity of bacterial taxa; therefore, we decided not to exclude REDgroups with very few members from the dataset. We also ran an additional random sampling analysis using the most populated REDgroups and confirmed the reproducibility of our results. Nonetheless, the remaining bias will only be eliminated with the inclusion of increased biodiversity in sequencing projects^17,20^.

Our analysis identified a plethora of unexplored taxonomic groups with substantial biosynthetic potential^9,10,48–50^. At the same time, it revealed that undiscovered biosynthetic diversity present in well-characterized NP producers. For example, multiple Proteobacteria taxa were identified among the top producers: *Pseudomonas*, *Pseudoalteromonas*, *Paracoccus*, *Serratia* among others. This is in accordance with the known biosynthetic potential of the Proteobacteria phylum^36^. Furthermore, we identified taxa that are less well represented in sequence databases as being potentially useful sources of secondary metabolites, including myxobacterial genera *Cystobacter*, *Melittangium*, *Archangium*, *Vitiosangium*, *Sorangium* and *Myxococcus^9,33,51^*, and *Chryseobacterium* and *Chryseobacterium_A^52^* from the Bacteroidota phylum. However, the most diverse groups of metabolites are predicted to be produced by actinobacterial strains of well-known and well-studied NP producers such as *Actinoplanes*, *Amycolatopsis*, *Micromonospora*, *Mycobacterium*, *Nocardia* and *Streptomyces^8,26,27,37^*. These bacteria produce most of the natural product antibiotics^26^ and our analysis confirms that recent analyses of biosynthetic novelty in the genomes of rare actinobacteria suggest that there is still much more natural product diversity to be discovered in this group as more diversified strains get sequenced^8,26,27,53^.

*Streptomyces* is a genus of the Actinobacteria phylum that contains some of the most complex bacteria that we know of, though by far not the most sequenced in our dataset (Supplementary Figure 5). These bacteria have been known as NP producers for a long time^37^, as single strains containing a high number of BGCs have been discovered, taking up to 10% of their genome^54^. However, members of other genera contain comparable absolute numbers of BGCs. This is the first time that a systematic comparison of the diversity of the encoded compounds within bacterial genera has been conducted, revealing how diverse *Streptomyces* are compared to all others^37^. The factors that cause this taxonomic group to stand out are not completely clear but probably related to their sophisticated lifestyle. Many observations suggest that NP biosynthesis drives speciation within the *Streptomyces* genus^8^. The exploration of factors that led to the rise of biosynthetic diversity in *Streptomyces* to such an impressive degree will be the subject of further investigations in the future.

Having the genomic capacity for the biosynthesis of secondary metabolites does not always herald the discovery of a novel chemistry^55,56^. Sometimes, the bacterium in question cannot be grown or BGCs are not expressed in laboratory conditions^26,48,50,55,56^. This issue is related to the complexity of BGCs; we have only just scratched the surface of their intricate regulation and connection to primary metabolism^5,48,55,57^. However, efforts to decode biosynthetic mechanisms for the activation of silent clusters need to be tailored to specific producer groups^26,27,56^, such as groups phylogenetically related to promising producers, e.g. members of the Pseudonocardiaceae family (REDgroups Amycolatopsis_RG1 & Kutzneria in Figure 4, these and more REDgroups in Supplementary Table 8), partly on the grounds that each phylum has unique diversity (Figure 2b).

Original approaches to the prioritization issue of NP research continue to emerge, fuelled by the advances in metagenomics and computational tools that enable the use of the biosynthetic potential of unculturable bacteria from environmental samples^58^. Furthermore, apart from the few metagenomic projects whose MAGs we incorporated in the first part of our analysis, there are multiple such projects publicly available, some of which have been the focus of NP studies^59^. Although the reconstruction of genomes from metagenomes remains a challenge^60^ and the assembly will often miss BGCs^61^, which has indirectly prevented their comparison to the cultured bacteria in the current project, metagenomics is proving a promising source of information on NPs and their producers^7,37,48,58,59^, as made apparent in the present investigation. We expect the effect of this field on NP research to become more evident in the following years.

The collection of microbial data from a large variety of habitats points to another interesting aspect, namely the relation between the biome of origin of the producers and the uniqueness of their biosynthetic diversity. Although this connection has been investigated to some extent^24,25,35,36,50^ drawing more definitive conclusions will require the use of a wider-scale dataset and the availability of more detailed and standardized metadata of producers’ genomes.

Our analysis provides a global overview of diverse known and promising understudied NP-producing taxa. We expect this to greatly help overcome one of the main bottlenecks of Natural Product discovery: the prioritization of producers for research^58^.

## Online Methods

### BGC data set

We obtained 170,585 complete and draft bacterial genomes (Table 1) from RefSeq^13^ on 27 March 2020. Furthermore, a dataset of 47,098 MAGs was included in the first part of the analysis (see Results: Biosynthetic diversity of the bacterial kingdom). For the rest of the study, we used only 161,290 RefSeq bacterial genomes whose taxonomic classification up to the species level was known (Table 1). All genomes were analyzed with antiSMASH (version 5)^12^, which identified their BGCs (Supplementary Table 1). The entirety of the MIBiG^23^ database (accessed on 27 March 2020) was included in parts of our analysis (their IDs can be found in Supplementary Table 6).

### Taxonomic classification

Due to multiple indications regarding a lack of accuracy of NCBI’s taxonomic classification of bacterial genomes^43–46^, we chose to use the Genome Taxonomy Database (GTDB^14^) instead. The bacterial tree of 120 concatenated proteins (GTDB release 89), as well as the classifications of organisms up to the species level, were included in the analysis.

### Quantification of biosynthetic diversity with BiG-SLiCE

For a bacterium to be regarded as biosynthetically diverse, we considered not the number of BGCs important, but rather how different these BGCs are to each other. In order to quantify this diversity, we analyzed all BGCs with the new BiG-SLiCE tool^21^, which groups similar clusters into Gene Cluster Families (GCFs). However, the first version of this tool has an inherent bias towards multi-protein families BGCs, producing uneven coverage between BGCs of different classes (i.e., due to their lack of biosynthetic domain diversity, all lanthipeptide BGCs may be grouped together using the Euclidean threshold of T=900, which in contrast is ideal for clustering Type-I Polyketide BGCs). To alleviate this issue and provide a fair measurement of biosynthetic diversity between the taxa, we modified the original distance measurement by normalizing the BGC features under L^2-norm, which will produce a cosine-like distance when processed by the Euclidean-based BIRCH algorithm. This usage of cosine-like distance will virtually balance the measured distance between BGCs with “high” and “low” feature counts (Supplementary Figure 6a), in the end providing an improved clustering performance when measured using the reference data of manually-curated MIBiG GCFs (Supplementary Figure 6b).

The GTDB^14^ (release 89) bacterial tree was pruned so that it included only the organisms that are part of our dataset. Then, having both the taxonomic classification of all bacteria, as well as how many GCFs their BGCs group into, the pruned GTDB tree was decorated with #GCFs values at each node. This allowed for the evaluation of the biosynthetic diversity of any clade, including the main taxonomic ranks. To pick a single threshold for subsequent taxonomy richness analysis, we compared BiG-SLiCE results on 947 MIBiG BGCs versus the compound-based clustering provided by the NPAtlas database^11^ (Supplementary Figure 2). A final threshold of T=0.4 was chosen based on its similarity to NPAtlas’s compound clusters (V-score=0.9X, GCF counts difference=+XX).

### Quantification of biosynthetic diversity with clust-o-matic

We aimed to repeat and evaluate the reproducibility of the BGC-to-GCF quantification step of BiG-SLiCE with an alternative, independently derived algorithm. For that instead of grouping BGCs into GCFs based on biosynthetic domain diversity, we developed an algorithm that considers full core biosynthetic genes. Biosynthetic gene clusters that were detected in the input data by antiSMASH 5.1 were parsed to deliver core biosynthetic protein sequences. Those protein sequences were subjected to all-against-all multi-gene sequence similarity search with DIAMOND^62^ 2.0 using default settings. Only one best hit per query core gene per BGC was allowed divided by a total core protein length, resulting in the final pairwise BGC score always being within range of 0 to 1. Pairwise BGC similarity scores were used to build a distance matrix that was later subjected to agglomerative hierarchical clustering in python programming language (package scipy.cluster.hierarchy). The same process as described in the paragraph above (for BiG-SLiCE in that case) was performed for identification of the most suitable threshold for the clust-o-matic algorithm. The determined optimal threshold of 0.5 was then used to generate GCFs, which were then fed into the next steps in parallel to the original set of GCFs obtained from BiG-SLiCE.

### Biogeography Analysis

One^20^ of the MAGs datasets was accompanied by sufficient metadata that allowed for a study of a potential connection between biosynthetic diversity patterns and the biomes of origin of the corresponding MAGs. The GCFs for each ecosystem type were collected by combining information from Supplementary Tables 1, 2 of this project and from the Nayfach paper^20^ Supplementary Information. This led to the creation of Supplementary Table 7. Then, the largest occurring intersections were computed and visualised in Extended Data Figure 2 using the UpSet^63^ visualisation technique.

### Variance Analysis

In order to pinpoint the emergence of biosynthetic diversity, the within-taxon homogeneity was compared among the main taxonomic ranks. For each rank, the variance value was computed (with NumPy^64^) based on the #GCFs values of immediately lower-ranked taxa, as long as there were at least two such taxa. For example, a phylum that includes only one class in our dataset was omitted from this computation. But a phylum with two or more classes would be assigned a variance value computed from its classes’ #GCFs values. The distribution of these variance values was plotted for each rank in Figure 3a. We noticed a significant reduction in variance from the family to the genus rank, which was confirmed with an additional statistical test (Supplementary Figure 3, Supplementary Methods). A similar variance analysis was performed to compare genera and REDgroups (Supplementary Figure 4) but in this case variance was calculated based on the strains’ biosynthetic diversity.

### Definition of REDgroups

To study the biosynthetic diversity of genera, we attempted to achieve uniform taxa. The creators of GTDB used Relative Evolutionary Divergence (RED) for taxonomic rank normalization^14^; it is a metric that relies heavily on the branch length of a phylogenetic tree and is consequently dependent on the rooting. The GTDB developers provided us with a bacterial tree decorated with the average RED values of all plausible rootings at each node. Since GTDB accepts a range of RED values for each taxonomic rank placement^14^, we chose the median of GTDB genus RED values, namely 0.934, as a cutoff threshold. Any clade in the GTDB bacterial tree with an assigned RED value higher than the threshold was considered one group (Supplementary Figure 7) that we named “REDgroup”. For REDgroup naming conventions, see Supplementary Figure 7.

### Rarefaction analysis

The extrapolation of potential #GCFs values was achieved by conducting rarefaction analyses, by use of the iNEXT R package^65^. A GCF presence/absence table (GCF-by-strain matrix) was constructed for each group considered and was then used as “incidence-raw” data in the iNEXT main function, where 500 points were inter- or extrapolated with an endpoint of 5000 for the REDgroups, and of 1.6 million (about 8 times the number of strains in the Complete Dataset) in each group for the RefSeq analyses (where 2000 points were inter- or extrapolated). By default, the number of bootstrap replications is 50.

### Random sampling

In order to test whether the above methods (creation of REDgroups and the subsequent rarefaction analyses) overcome the inherent sequencing bias in our dataset, a random sampling technique was used. A reduced dataset was tested that included only those REDgroups containing at least 20 members. For each REDgroup, a sample of 20 genomes was randomly chosen (using the Python “random” module), while preserving the species diversity of the group. The latter was achieved by ensuring that genomes belonging to as many species as possible are included in each sample; if all species of a REDgroup were included but the genomes were fewer than 20, the remaining “spots” were distributed evenly among a random sample of the REDgroup’s species. One hundred iterations of this process were calculated for all REDgroups in this reduced dataset and rarefaction analyses were conducted for the random samples in each iteration. Finally, the average potential GCFs (pGCFs) value for each REDgroup from all iterations was calculated and reported in Supplementary Table 9.

### Identification of unknown producers

We investigated the genera included in the most promising REDgroups, to find out whether they include species that are producers of known compounds. Hence, the species names were cross-referenced with the species named as producers in the NPASS depository^38^ (accessed on 15 October 2020), taking care to match the GTDB-given names to the NCBI-given names that the database uses.

## Data availability

The datasets generated and analyzed during the current study are available in the following zenodo repository: https://doi.org/10.5281/zenodo.6365726.

## Code availability

The clust-o-matic code is available here: https://github.com/Helmholtz-HIPS

The modified BiG-SLiCE script (that accepts as input a regular BiG-SLiCE output folder, then outputs the GCF membership in a tsv file) is available both in our zenodo repository (file name: perform_l2norm_clustering.py) and under the following link: https://github.com/medema-group/bigslice/blob/master/misc/useful_scripts/perform_l2norm_clustering.py

## Supplementary information

### Supplementary Methods and Figures

Supplementary Methods and Supplementary Figures 1-7 with their legends are provided in an additional file.

**Supplementary Table 1**

Accession numbers, GTDB-based taxonomic information and BGC IDs of all genomes from all datasets used in the analysis. MAG datasets: mag_uba^19^, mag_humangut^17^, mag_chicken^16^, mag_bovine^15^, mag_ocean^18^, GEMS^20^. Available in the zenodo repository.

**Supplementary Table 2**

BGC to BiG-SLiCE GCF assignment and centroid distance for T=0.4 (which proved to be the most suitable threshold). Available in the zenodo repository.

**Supplementary Table 3**

BGC to BiG-SLiCE GCF assignment and centroid distance for T=0.5. Available in the zenodo repository.

**Supplementary Table 4**

BGC to BiG-SLiCE GCF assignment and centroid distance for T=0.6. Available in the zenodo repository.

**Supplementary Table 5**

BGC to BiG-SLiCE GCF assignment and centroid distance for T=0.7. Available in the zenodo repository.

**Supplementary Table 6**

BGC IDs, MiBIG IDs, producer GTDB-based taxonomic information and GCF assignment (for BiG-SLiCE T=0.4) for all MiBIG BGCs included in the creation of Figure 1C.

**Supplementary Table 7**

Biogeography analysis of the Nayfach MAGs dataset^20^. Number of genomes, BGCs, GCFs and unique GCFs per ecosystem type (as defined in the corresponding paper^20^).

**Supplementary Table 8**

REDgroup full metadata: Node IDs (can be used in the exploration of the tree in Extended Data 3), labels, number of members, number of BGCs, number of GCFs and potential GCFs (pGCFs) as defined by BiG-SLiCE (T=0.4) and clust-o-matic (T=0.5), GTDB taxonomic information and number of products in the NPASS database whose producer is a member of the REDgroup (NPASS_hits).

**Supplementary Table 9**

Comparison of random sampling analysis results to original results. The table includes: Node IDs, labels, number of members, number of BGCs, number of GCFs and potential GCFs (pGCFs) as defined by BiG-SLiCE (T=0.4), the original ranking based on the pGCFs, the average pGCFs from all random sampling iterations, the ranking based on the random sampling and GTDB taxonomic information.

## Acknowledgements

A.G. is grateful for the support of the Deutsche Forschungsgemeinschaft (DFG; Project ID # 398967434-TRR 261). N.Zi. is supported by the German Center for Infection Research (DZIF) (TTU 09.716). M.H.M. is supported by an European Research Council Starting Grant 948770-DECIPHER. S.K. was supported by the Graduate School for Experimental Plant Sciences (EPS) of Wageningen University. Work in the lab of R.M. is supported by BMBF (16GW0243), DFG and DZIF (807-5-8-0982600). A.G. and N.Zi. thank the Deutsche Forschungsgemeinschaft (DFG, German Research Foundation) under Germany’s Excellence Strategy – EXC 2124 – 390838134 for the infrastructural support. A.G. thanks M. Direnc Mungan for valued discussions on optimizing the analysis, as well as Caner Bagci for his imaginative suggestion on dealing with large data. We also thank Dr. Libera do Presti for invaluable comments on the manuscript.

## Author contributions

A.G., S.A.K., N.Za. and D.K. have performed the analysis. S.A.K. and N.Za. have contributed analysis tools. A.G., D.K., R.M., M.H.M. and N.Zi. have written the paper. All authors have contributed to the conception and design of the analysis. All authors have read and agreed to the published version of the manuscript.

## Competing interests

M.H.M. is a co-founder of Design Pharmaceuticals and a member of the scientific advisory board of Hexagon Bio. The other authors declare no competing interests.

## Notes

https://doi.org/10.5281/zenodo.6365726

### Summary of Updates

Small changes have been made in various locations to improve readability and provide additional explanation where needed. The list of references has been reduced. The DOI of the zenodo repository has been updated to the latest version.

