## Supplementary Information for "Compendium of secondary metabolite biosynthetic diversity encoded in bacterial genomes"

### Supplementary Methods

##### *Statistical analysis of GCF profile similarity*

We analyzed the similarity of genomic BGC content as a function of phylogenetic relatedness, on the basis of GCF distributions from 163,269 strains spanning 1,707 bacterial genera. For this purpose the occurrence pattern of 41,870 GCFs was converted into a matrix comprising 163,269 binary profiles and pairwise similarity was determined using the bitvector cosine similarity measure. We found that BGC profile similarity clearly decreases as taxonomic distance increases (Supplementary Figure 3). Apart from the clearly visible shift, the means of similarity distributions for taxonomic ranks were also statistically compared using Wilcoxon-Mann-Whitney tests, to yield p values close to 0.

### Supplementary Figures

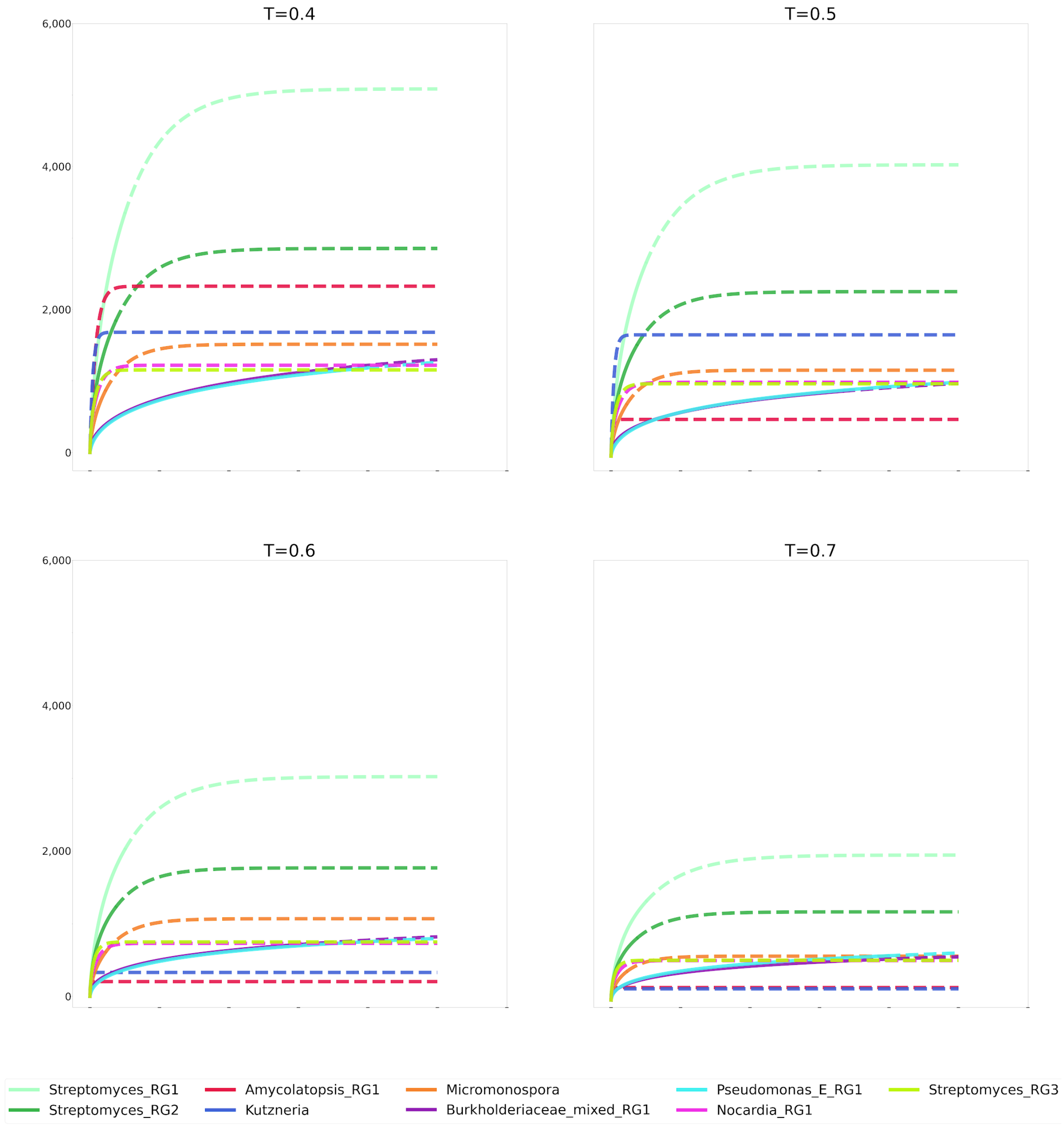


**Supplementary Figure 1**: Rarefaction curves (see Methods: rarefaction curves) of promising REDgroups (see Methods: REDgroups definition) in different BiG-SLiCE thresholds (extrapolated up to 5,000 sequences). The solid lines represent interpolated and actual data, while the dotted lines represent extrapolated data. The absolute number of potential Gene Cluster Families (GCFs, as defined by BiG-SLiCE) changes from threshold to threshold, but the general tendencies (most to least promising group) remain the same.

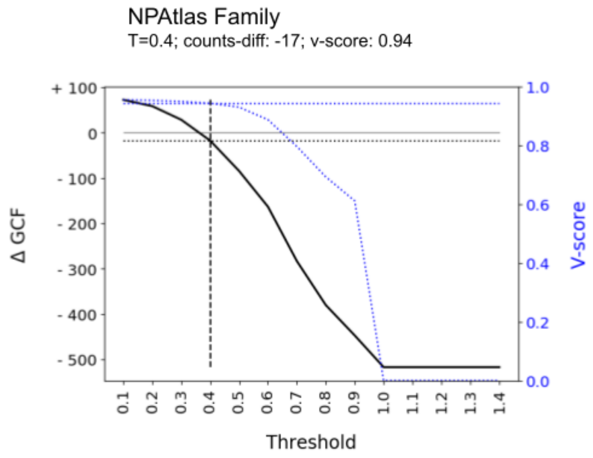


**Supplementary Figure 2: mapping the grouping of 947 known BGCs with link to NPAtlas compounds.** Clustering agreements between NPAtlas families of the compounds and BiG-SLiCE GCFs are compared across different thresholds, as measured by the v-score (dashed blue line) and differences in the total number of GCFs (solid black line). In the end, T=0.4 were picked as the threshold to use for our analysis, which shows the best agreement with the NPAtlas family assignments (v-score=0.94, ΔGCF=-17).

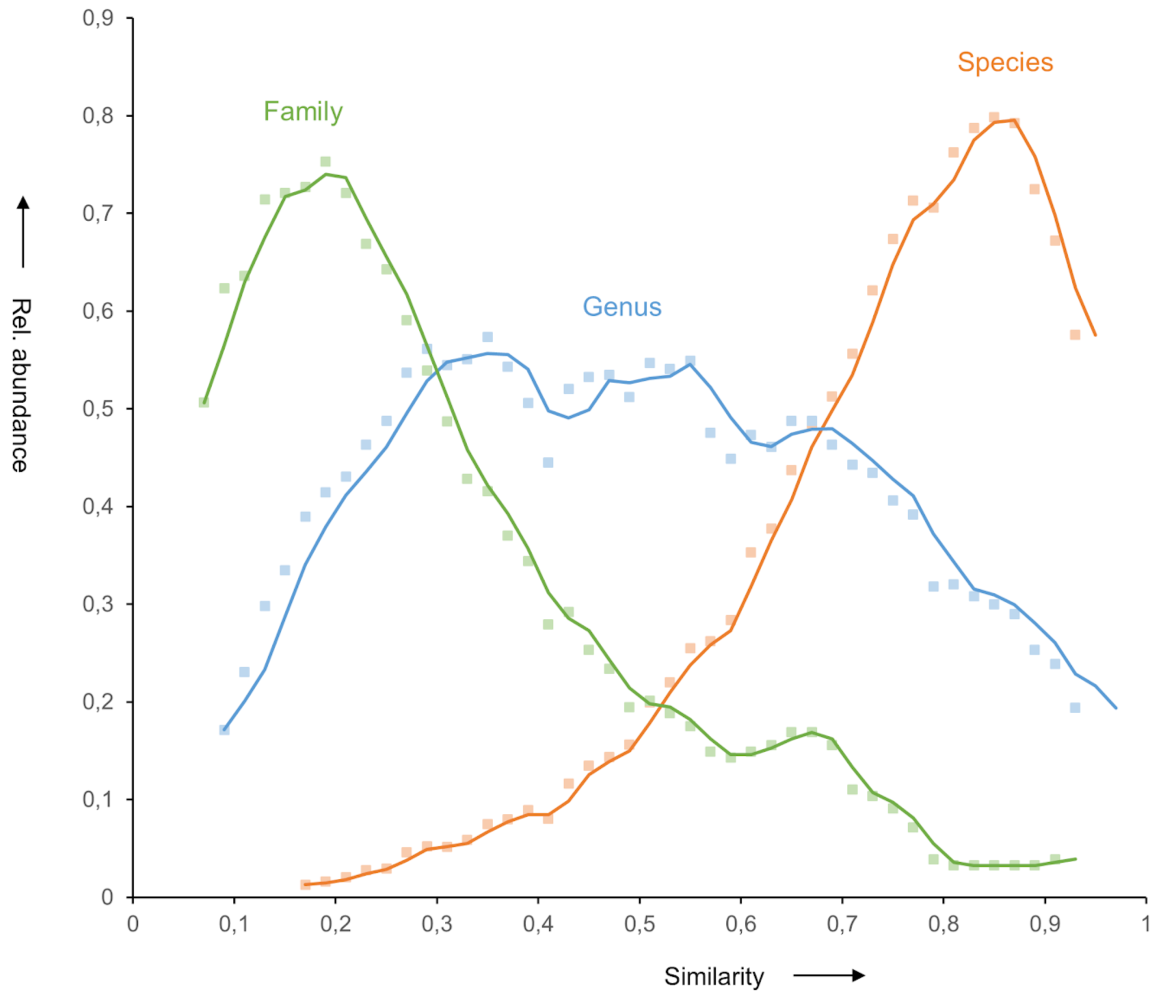


**Supplementary Figure 3: BGC content similarity in relation to taxonomic distance.** Histogram plots depict binned distributions for genomic BGC content similarity within and between varying taxonomic ranks, where 1 on x axis scale means strains contain highly similar GCFs (as defined by BiG-SLiCE with T=0.4). Similarity distributions were calculated using GCF profiles from: various strains within their respective species (orange); strains belonging to different species but within their respective genus (blue); strains belonging to different genera but within their respective family (green). Dots represent relative abundance values binned in 2% intervals, curves were smoothed by moving-average with a period length of 2. See also Supplementary Methods.

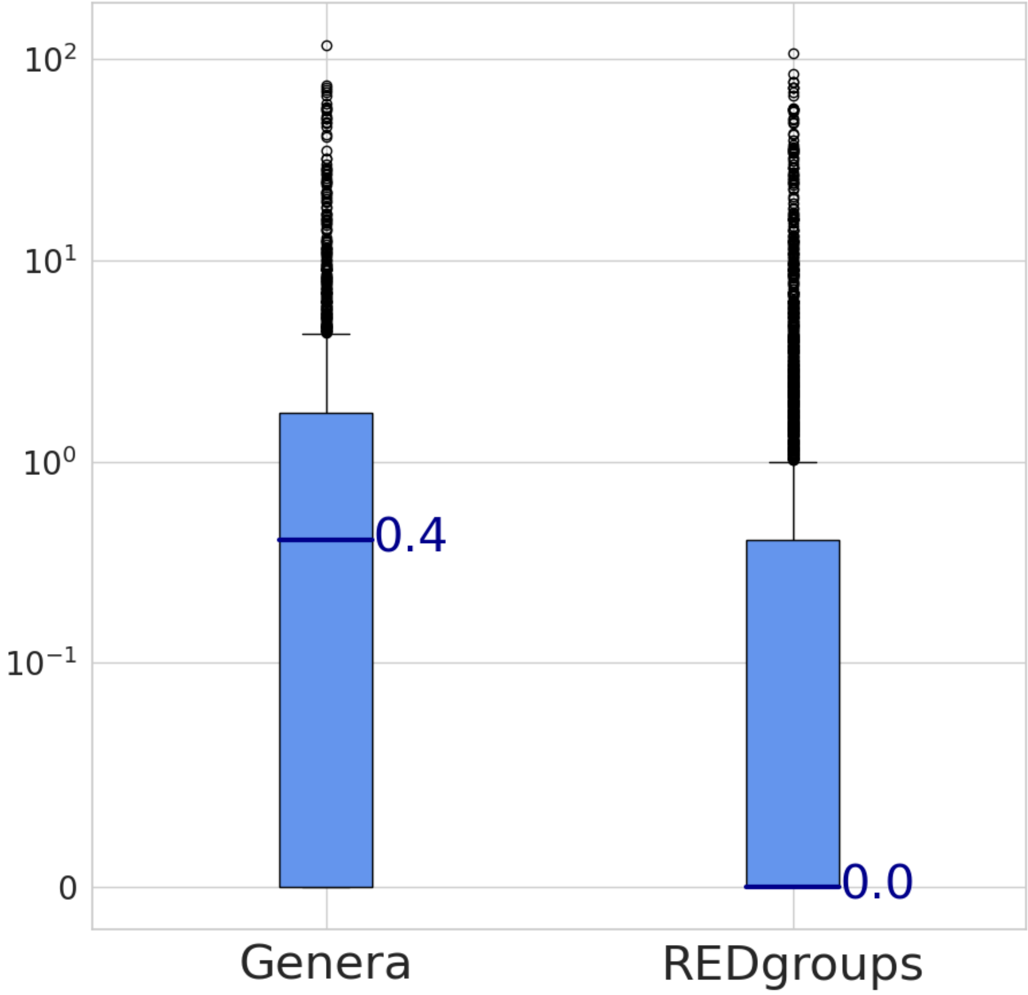


**Supplementary Figure 4: Comparison of biosynthetic diversity variance between Genera and REDgroups.** Each boxplot represents the dispersion of variance values of the genera and REDgroups classifications, computed from the number of Gene Cluster Families (GCFs as defined by BiG-SLiCE at T=0.4) of the strains belonging to each group. The boxplots' center line represents the median value; the box limits represent the upper and lower quartiles. Whiskers represent a 1.5x interquartile range. Points outside of the whiskers are outliers. Sample sizes are: Genera n=1,607, REDgroups n=3,779. There is significant difference in dispersion of variance values between the genera and the REDgroups (verified using a Wilcoxon-Mann-Whitney test with a p-value very close to zero), indicating that biosynthetic diversity is more uniformly distributed in the latter.

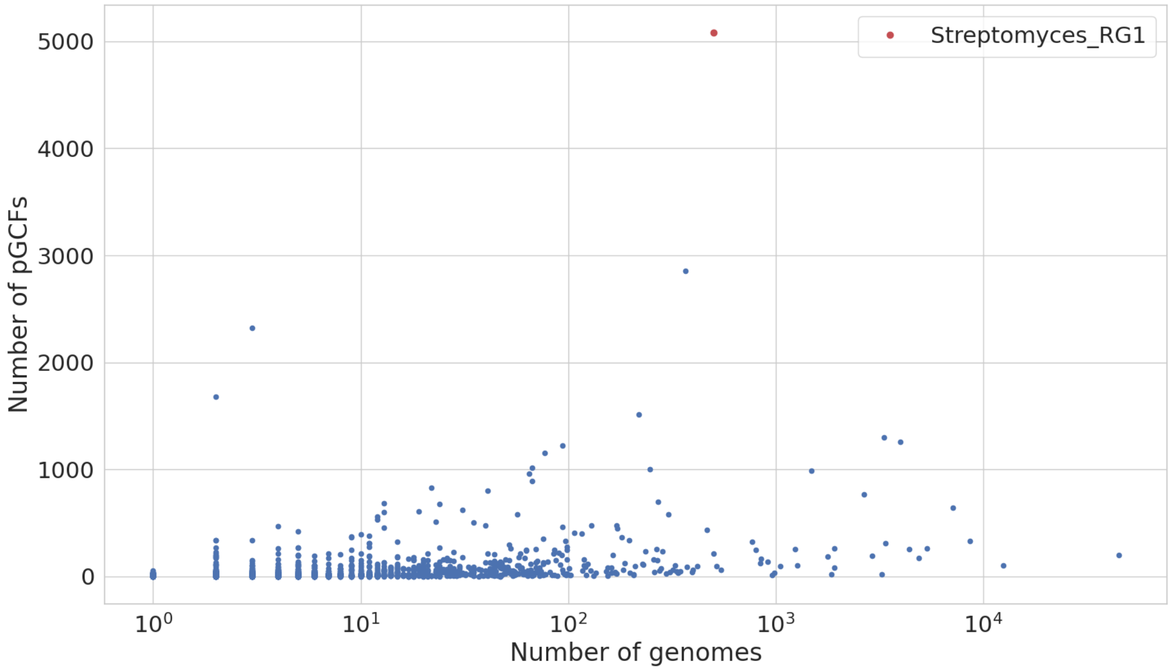


**Supplementary Figure 5: Examining the correlation between number of genomes and biosynthetic potential in REDgroups.** For each REDgroup (see Methods), the number of genomes was plotted against the number of potential GCFs (pGCFs), as extrapolated by rarefaction analyses from GCFs defined by BiG-SLiCE (T=0.4). The x axis is in logarithmic scale. The most promising REDgroup, Streptomyces_RG1, is marked with red and labeled; it is by far not the most sequenced group but its potential is unparalleled. There does not appear to be a clear correlation between the raw number of genomes sequenced and their predicted biosynthetic diversity, as calculated with the methods described in this study.

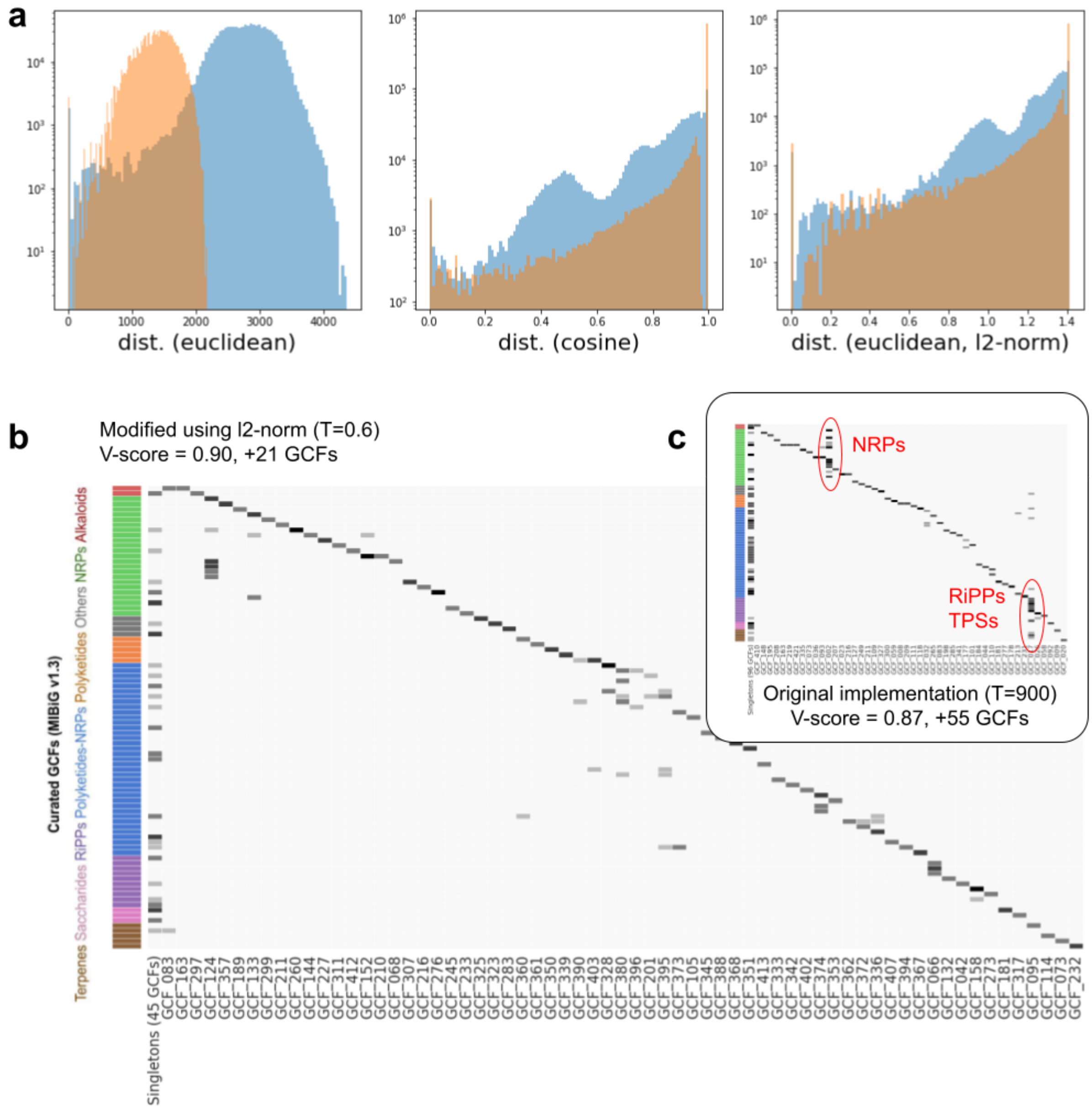


**Supplementary Figure 6.** **a**: Histograms comparing the pairwise-distance distribution across 1,000+1,000 randomly picked BGCs in the dataset, half were selected with "low" feature counts (< 30, orange-colored bars) while the other half having "high" feature counts (>= 30, blue-colored bars). It was pretty clear that the usage of euclidean distance on the unmodified BiG-SLiCE features resulted in a bias in distance calculations, which in turn resulted in an uneven clustering across different BGC classes (e.g., NRPs, RiPPs and TPSs were often over-clustered in the original BiG-SLiCE implementation), as shown in panel **c**. The usage of cosine distances greatly alleviates this issue, however it is not supported by the BIRCH clustering algorithm used by BiG-SLiCE. By converting the original features into an l2-normalized feature matrix, we showed that we can achieve similar results while still using euclidean-based distances, in the end resulting in a much more "balanced" clustering across the board (panel **b**).

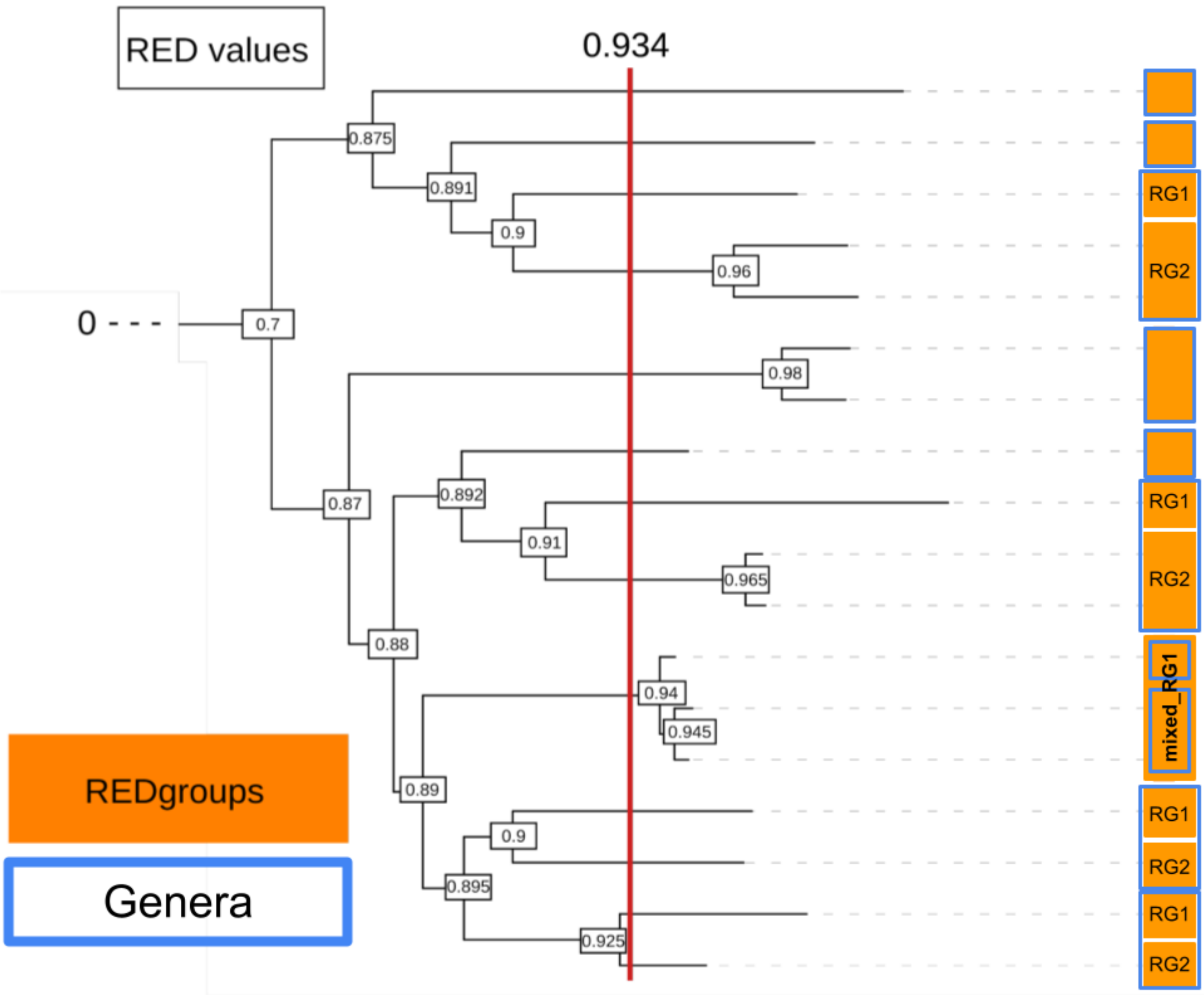


**Supplementary Figure 7: Creation of REDgroups (mock example)**. The GTDB (Genome Taxonomy DataBase) bacterial tree was decorated with RED (Relative Evolutionary Divergence) values at each node (black outlined rectangles). Every clade whose earliest node has a RED value above the threshold of 0.934 (red line) was considered a REDgroup (orange filled rectangles). REDgroups that correspond exactly to a genus (blue outlined rectangles) retain the genus name, but are not written in italics. REDgroups that include only part of a genus receive a suffix “RG” plus a number (so a genus “Alpha” could be separated to REDgroups: Alpha_RG1, Alpha_RG2, Alpha_RG3). When a REDgroup encompasses more than one genus it is named after the family and receives the suffix “mixed_RG” plus a number (so a family “Beta” could have 3 REDgroups each corresponding to a genus - or part of one - and another REDgroup with the rest of the family’s genera that is called “Beta_mixed_RG1”).
