## Extended Data for "Compendium of secondary metabolite biosynthetic diversity encoded in bacterial genomes"

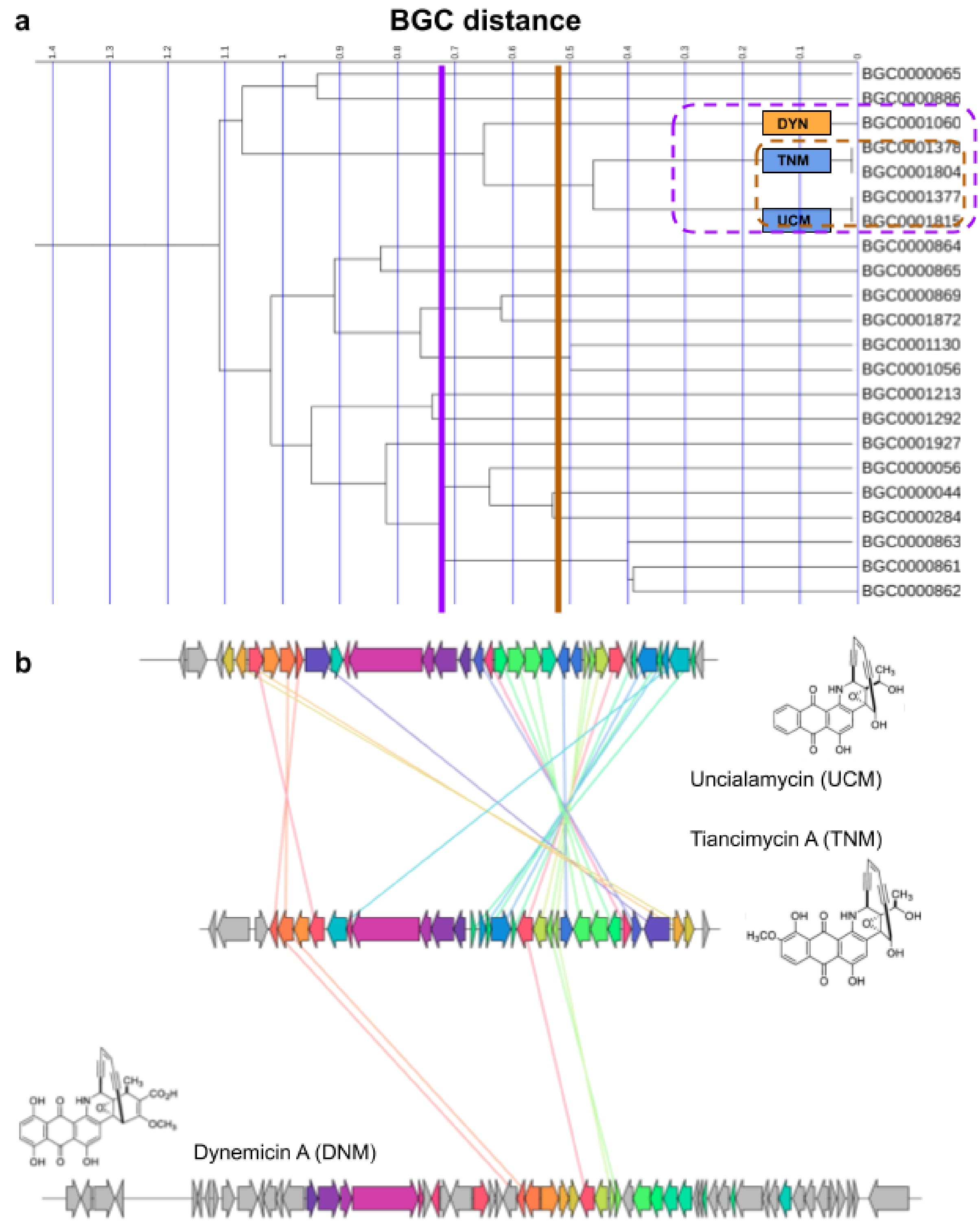


**Extended Data Figure 1**: Illustrating the correlation between BGC clustering thresholds and the grouping of their pathway products. a) a snippet of a complete-linkage hierarchical dendrogram constructed by doing a pairwise distance comparison of L2-normalized BGC features within the MIBiG dataset, highlighting the grouping of BGCs for the enediynes Uncialamycin (UCM) and Tiancimycin (TNM) under the threshold T=0.5, and further grouping with another related enediyne BGC, Dynemicin (DNM) under the looser threshold of T=0.7. b) Comparative genes analysis generated using the clinker tool[92] v0.0.23 shows how UCM and TNM BGCs are much more similar to each other than to DNM (same-colored genes indicate <70% amino acid similarity, while colored edges indicate <50% amino acid similarity), which is consistent with the structural diversity of their compounds (pictured).

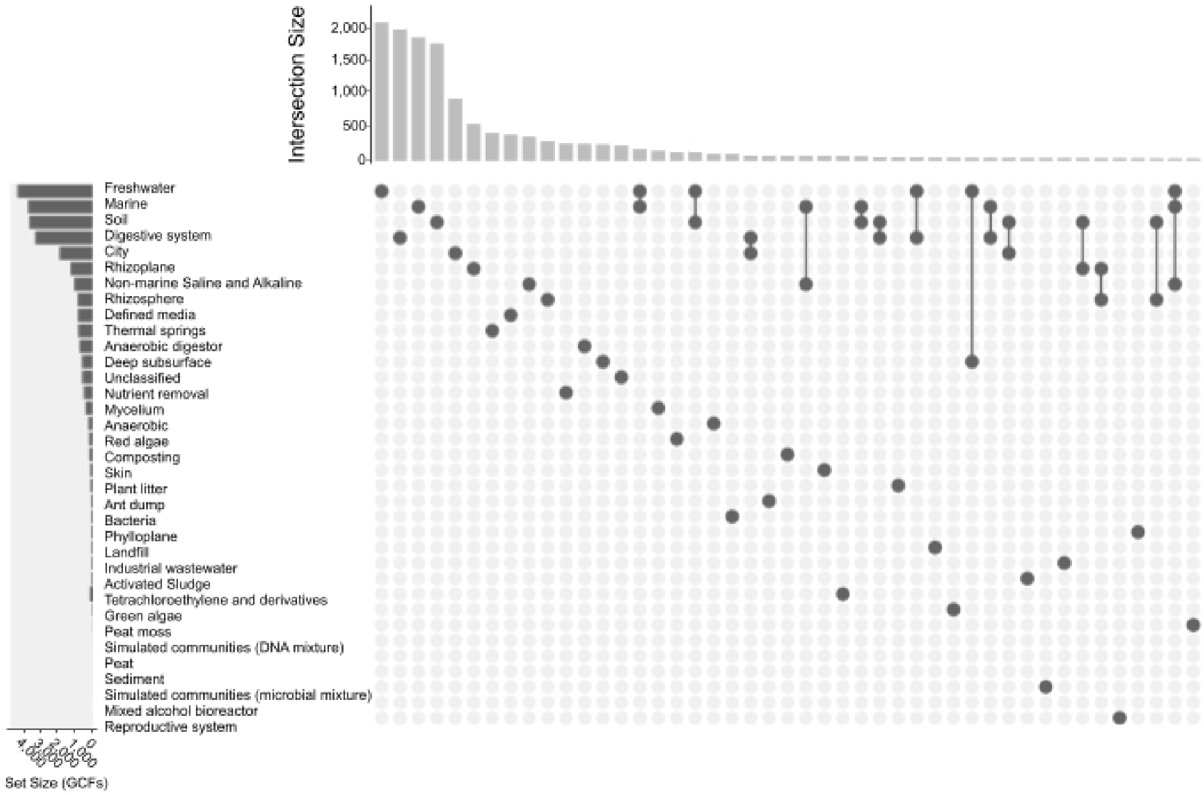


**Extended Data Figure 2**: Intersections and distribution of biosynthetic diversity values among different ecosystem types. The bar plot on the left depicts the number of Gene Cluster Families (GCFs as defined by BiG-SLiCE with T=0.4) found in each biome type. The bar plot on top shows the size (number of GCFs) of each intersection. Which sets (biome types) are included in each intersection can be seen in the matrix below the bar plot, where the dark dots pinpoint included sets. If more than one set is part of an intersection, connecting lines are drawn for better visibility. The data presented in this graph come only from the MAGs in the GEMS dataset (see Supplementary Table 1), which was the only one with sufficient metadata. Only the top 63 most sizable intersections are depicted here, and only the 35 ecosystem types (with the most GCFs out of the 63) that were part of them are shown on the left. The data indicate that there is barely any overlap between the ecosystem types; most GCFs (74.43 %) are specific to a single biome (a complete overview of unique GCFs per ecosystem type can be found in Supplementary Table 7), while the largest intersection (the one including most habitats - not visible in this Figure) includes 50 of the 63 ecosystem types.

**Extended Data Figure 3**: Overview of actual and potential biosynthetic diversity of bacterial kingdom, compared at REDgroup level. Extended Data Figure 3 is interactive and can be accessed online on iTOL: [https://itol.embl.de/shared/1B6W5n9MixSdJ](https://itol.embl.de/shared/1B6W5n9MixSdJ). GTDB bacterial tree up to REDgroup level (for more details see Methods - REDgroup definition), colour coded by phylum, decorated with barplots of actual (orange) and potential (purple) Gene Cluster Families (GCFs) as defined by BiG-SLiCE (T=0.4). Potential GCFs were computed by rarefaction analyses (for more details see Results - Well known and less popular taxa as sources of biosynthetic diversity). REDgroups names are displayed around the tree as leaf node labels; hovering over them provides further taxonomic information (for full REDgroup metadata see Supplementary Table 1). Phyla known to be enriched in NP producers are immediately visible (Actinobacteriota, Protobacteriota), with the most promising groups coming from the Actinobacteriota phylum (the highest peak belongs to a REDgroup containing Streptomyces strains). Simultaneously, within the underexplored phyla, there seems to be significant biosynthetic diversity and potential. This Figure is meant to be explored by zooming in and out, searching for keywords and visualizing different kinds of information by switching between Tree Views. Any other attempt at modification (e.g. turning datasets on and off) may result in an unreadable graph.

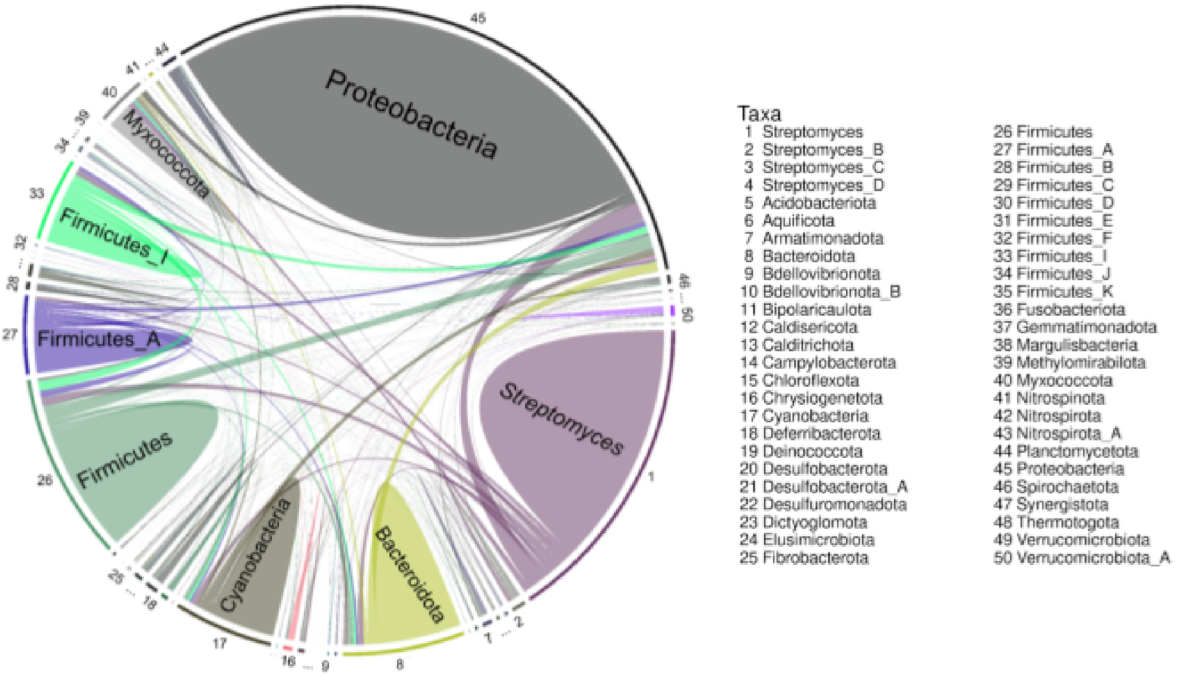


**Extended Data Figure 4**: Unique diversity in the known producer *Streptomyces.* Unique GCFs, as defined by BiG-SLICE (T=0.4), of bacterial phyla and Streptomyces (solid shapes) and pairwise overlaps of phyla - phyla and phyla - Streptomyces (ribbons). Each taxon has a distinct colour. The genus Streptomyces (1) appears to have a very high amount of unique GCFs comparable to entire phyla, such as Proteobacteria (43).

**Extended Data Table 1**: Information on the most biosynthetically promising REDgroups. Information on the most biosynthetically promising REDgroups (BiG-SLiCE T=0.4).

| REDgroup | #members | #BGCs | #GCFs | #pGCFs |
|---|---|---|---|---|
| Streptomyces_RG1 | 501 | 16,381 | 3,339 | 5,084 |
| Streptomyces_RG2 | 368 | 12,104 | 1,831 | 2,853 |
| Amycolatopsis_RG1 | 3 | 84 | 83 | 2,324 |
| Kutzneria | 2 | 82 | 81 | 1,681 |
| Micromonospora | 219 | 5,015 | 855 | 1,513 |
| Burkholderiaceae_mixed_RG1 | 3,331 | 68,547 | 1,153 | 1,297 |
| Pseudomonas_E_RG1 | 3,980 | 39,055 | 1,181 | 1,260 |
| Nocardia_RG1 | 94 | 3,035 | 777 | 1,220 |
| Streptomyces_RG3 | 77 | 2,352 | 766 | 1,154 |
| Amycolatopsis_RG2 | 67 | 2,207 | 645 | 1,019 |
